# Affective flexibility without perceptual awareness

**DOI:** 10.1101/505545

**Authors:** Philipp Homan, H. Lee Lau, Ifat Levy, Candace M. Raio, Dominik R. Bach, David Carmel, Daniela Schiller

## Abstract

In an ever-changing environment, survival depends on learning which stimuli represent threat, and also on updating such associations when circumstances shift. Humans can acquire physiological responses to threat-associated stimuli even when they are unaware of them, but the role of awareness in *updating* threat contingencies remains unknown. This complex process – generating novel responses while simultaneously suppressing learned ones — relies on distinct neural mechanisms from initial learning, and has only been shown with awareness. Can it occur unconsciously? Here we show that it can. Participants underwent classical threat conditioning to visual stimuli that were suppressed from their awareness. One of two images was paired with an electric shock; halfway through the experiment, contingencies were reversed and the shock was paired with the other image. We found that physiological responses reflected changes in stimulus-threat pairings independently of stimulus awareness, demonstrating the sophistication of unconscious affective flexibility.

## Introduction

Flexible responses to environmental threats are essential for adaptive behavior. Cues that predict threat constantly change - new threats may arise while old ones cease to pose a risk. When consciously perceiving such cues, we are able to flexibly update and shift threat responses from one cue to another (1–3). But can we update our reaction to stimuli that predict danger when we are not aware of them? It is known that threat-conditioned stimuli that are perceived without awareness can still elicit defensive physiological reactions (4–7), and that new threat associations can be formed through classical conditioning even without any awareness of the conditioned stimuli (8–10). Updating threat associations when contingencies change, however, is an entirely different matter: it involves a complex process of creating novel responses while simultaneously suppressing acquired ones. To date, such updating has only been shown in humans who were aware of the stimuli (2), and in animals under conditions where stimuli were fully available for perceptual processing (11); these studies have shown, furthermore, that the neural substrates of threat updating differ from those of the initial learning. It is thus unknown whether the sophisticated re-evaluation involved in such affective flexibility requires awareness, or can be accomplished without it. Here we show that it can, and furthermore, that stimulus awareness does not seem to play a substantial role in such affective flexibility.

To examine this, we employed the reversal paradigm, a laboratory model that requires flexible updating of threat contingencies (2). In an initial acquisition phase, participants encounter two conditioned stimuli (CSs) and learn that only one of them predicts an electric shock. Halfway through the experiment, with no warning, these contingencies flip, initiating the reversal phase: Participants must flexibly learn that the formerly safe CS now predicts the shock and that the old one no longer does. To assess learning, participants’ physiological arousal is recorded throughout the experiment, typically (and here) by measuring their skin conductance responses. Appropriate response reversal requires a sophisticated form of updating, in that one must learn to respond to a cue that now predicts threat while simultaneously inhibiting responses to the previously threatening cue that is now safe.

To see whether reversal of conditioned threat requires awareness, we had a large group of participants (*N* = 86) undergo reversal learning with the CSs suppressed from awareness by continuous flash suppression (CFS), a technique commonly used to examine unconscious perception (10, 12–14): The CSs were visual images presented monocularly, while the other eye was shown a high-contrast, dynamic image (the CFS mask) at the corresponding retinal location (See Figure 1 for a description of the design and procedure).

**Figure 1:**
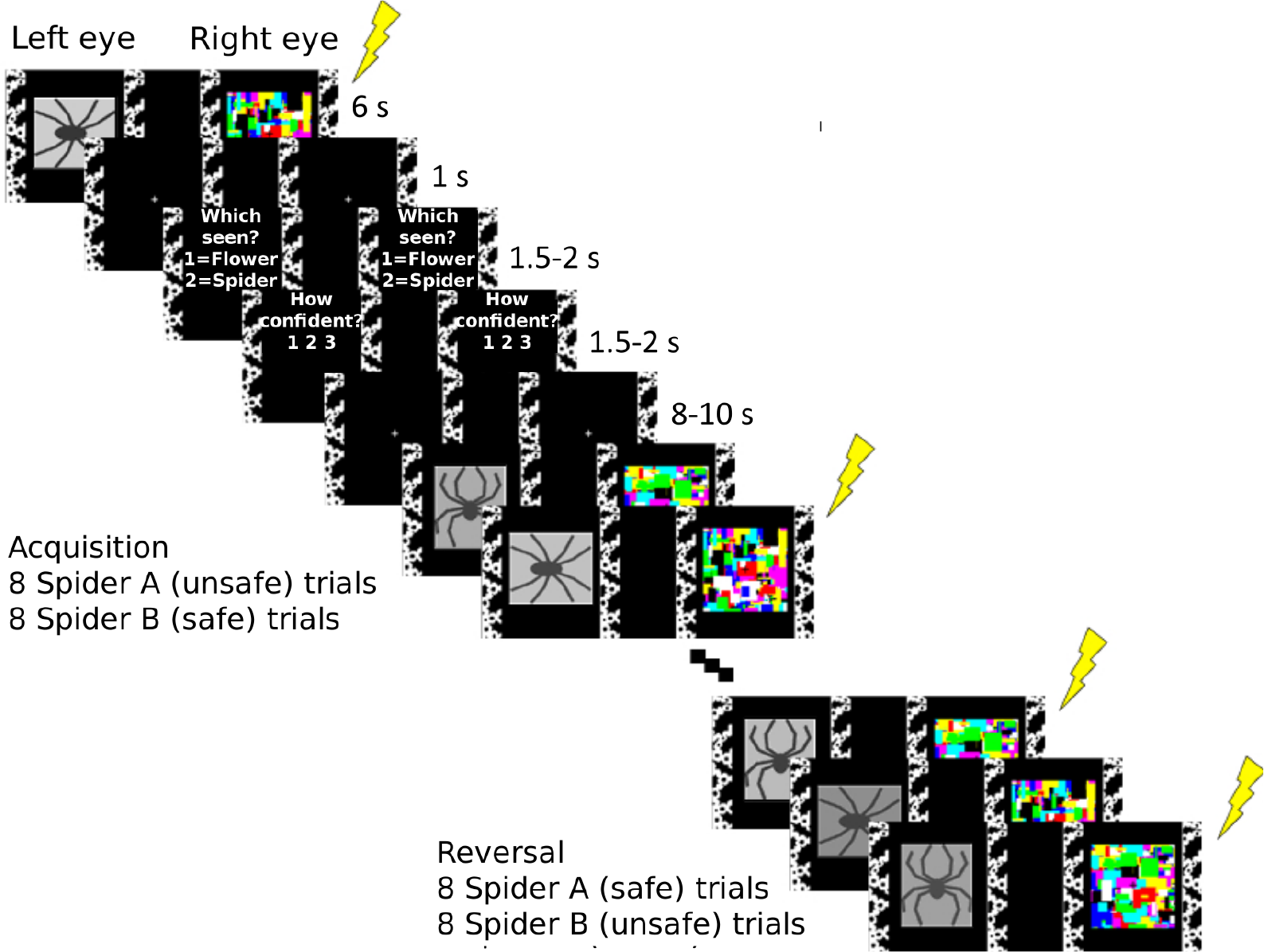
Schematic description of experimental design and procedure. In each trial of the acquisition phase, participants were presented with one of two stimuli (schematic pictures of spiders, presented monocularly for 6 sec and suppressed from awareness by a CFS mask shown to the other eye). One image (spider A) always terminated with a mild electric shock to the wrist, whereas the other (spider B) never did. Halfway through the experiment, with no warning, the contingencies flipped and the reversal phase began: the formerly safe stimulus (spider B) now predicted the shock, and the old threat-associated one (spider A) was now safe. Each spider was shown 8 times in each phase. Trial order was pseudorandomized (see Materials and Methods) and spider identity (A and B) was counterbalanced across participants. To assess the success of the awareness manipulation, participants answered the questions “Which seen?” (1=flower, 2=spider) and “How confident?” (1=guess to 3=sure), presented binocularly (1.5 - 2 s each), beginning 1 s after the offset of every CS, and followed by an 8-10 s inter-trial interval (the questions are only shown here for the first depicted trial, but were repeated in all trials). Participants who underwent the same procedure without CFS were shown identical CSs, but the CFS mask was absent.

CFS can suppress images from awareness for several seconds. However, it is also known that its effectiveness may vary across trials and individuals, and the suppressed stimulus may “break through” the suppression (15). Over the last decade, a growing body of work has raised concerns that the standard approach - removing from analysis data (participants and trials) in which breakthrough had occurred - may bias the findings (16, 17; See Supplementary Methods for further details of these issues.) Here, we adopt a number of methodological approaches to ensure our results are robust to these potential concerns.

Specifically, we remove no data and instead incorporate individual levels of reported stimulus awareness, as well as response patterns that might reflect residual awareness, into a regression model accounting for physiological responses. The model also adjusts for baseline anxiety (which has been previously shown to correlate with unconscious learning; (10)). Additionally, we use a Bayesian approach to establish that a model in which participants were updating their learning provides a better account for the findings than a model in which they were simply (and independently of the stimulus) predicting the probability of a shock on the next trial (18). Finally, in order to verify that our procedure is able to induce reversal learning when participants are aware of the stimuli (so that if no learning or updating were found under CFS, we could rule out the possibility this may be simply due to an ineffective procedure) we ran a no-CFS group (*N* = 12), in which participants also viewed the CSs monocularly (as the CFS group did), but were aware of them as no CFS masks were presented to their other eye.

We hypothesized that physiological responses to threat can be flexibly reversed without perceptual awareness. We find that reversal indeed occurs independently of CS awareness, and that there is strong evidence for the reversal of threat learning even in its complete absence.

## Results

### Overall assessment of physiological reversal learning

To assess the physiological arousal evoked by CSs, we used a model-based approach (19) to estimate the amplitude of anticipatory sudomotor nerve activity (SNA) from skin conductance data recorded during stimulus presentation. A variational Bayes approximation was employed to invert a forward model that describes how hidden SNA translates into observable SCRs (see Materials and Methods). Previous work has shown that this approach is more sensitive than conventional SCR peak-to-peak analysis (19–21). Figure 2A shows the time course of evoked SNA to Spiders A and B, separately for the CFS and no-CFS groups. In both groups, responses to Spider A relative to Spider B were larger during the acquisition phase and smaller during the reversal phase. To quantify the magnitude of physiological reversal learning, we calculated a reversal learning index for each participant (see Materials and Methods). The reversal learning index was positive and significantly greater than zero for both the CFS and no-CFS groups (Figure 2B). A linear mixed model (see Materials and Methods for details) revealed a significant interaction of stage and spider in both groups (CFS: β = 0.27, *t* (2935) = 4.23, *P* = < 0.001; no-CFS: β = 1.23, *t* (2935) = 7.29, *P* = < 0.001). Note that a significant interaction is formally equivalent to a significant reversal learning index. On its own, however, it simply reveals a difference in the comparative magnitude of responses to the two CSs across the two halves of the experiment; follow-up tests show that this difference is indeed due to reversal: Spider A evoked greater responses than Spider B in the acquisition phase (CFS: *t* (341.9) = 3.0, *P* = 0.003; no-CFS: *t* (201.1) = 4.6, *P* < 0.001) and the pattern was reversed in the reversal phase (CFS: *t* (341.9) = 2.8, *P* = 0.005; no-CFS: *t* (341.9) = 3.6, *P* = 0.0003). These results indicate that reversal learning was evident in both groups. Although Figure 2 shows that it was more pronounced in the no-CFS group, we note that this difference is not straightforwardly interpretable because the no-CFS group (a control, intended to rule out an ineffective manipulation if no effect was found for the CFS group) was substantially smaller; furthermore, as addressed in detail below, suppression from awareness was very heterogenous in the CFS group.

**Figure 2:**
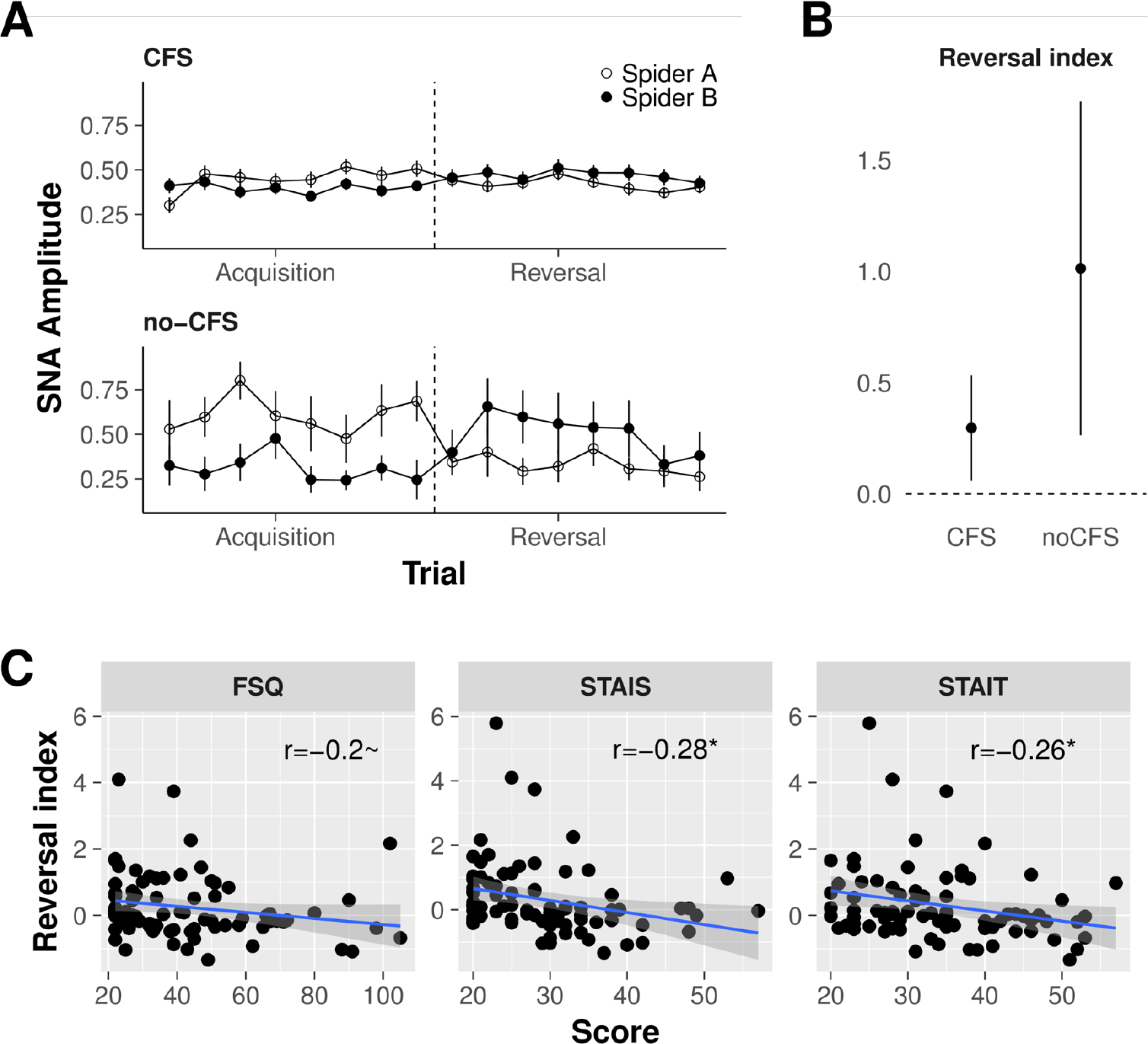
Physiological reversal learning. **A. Time courses reveal reversal of threat responses with and without continuous flash suppression**. Data points represent trial-wise mean responses to spider A (the CS+ during acquisition) and spider B (the CS- during acquisition). Both groups showed reversal learning, as indicated by greater responses to Spider A during the acquisition phase and greater responses to Spider B during the reversal phase. Error bars represent standard errors. **B. Mean reversal learning index for each group**. Error bars represent 95% confidence intervals, indicating that the interaction of stage and stimulus and thus the magnitude of reversal learning in both groups was significantly greater than zero. **C. Heightened anxiety is associated with impaired reversal learning under CFS**. A negative correlation between baseline anxiety measures and the strength of threat reversal learning is evident for state and trait anxiety. Blue lines show linear fits of each score to the reversal index, and ribbons around lines indicate bootstrapped 95% confidence intervals around the estimate. *Abbreviations*: STAIS/STAIT, state/trait anxiety subscale of the Spielberger State-Trait Anxiety Inventory; FSQ, Fear of Spider Questionnaire, ~, *P* < .1; *, *P* < .05.

As previous work has found a negative association between anxiety and threat acquisition with and without awareness (10), we also calculated correlations between the CFS group’s baseline anxiety measures (STAIT, STAIS, FSQ) and the reversal learning index. Overall, reversal learning decreased significantly with increasing levels of state and trait anxiety, and to a lesser but non-significant extent for spider phobia (Figure 2C).

### Reversal learning and perceptual awareness

The CFS manipulation reduced awareness of the CSs; as expected, however, it was differentially effective in doing so across participants, precluding an overall conclusion that all learning under CFS happened non-consciously. The CFS group showed significantly lower accuracy in response to the “which seen?” question (M = 0.46, SD = 0.29) compared to the no-CFS group (M = 0.86, SD = 0.16; *t* (22.77) = -7.24, *P* < 0.001), and accuracy in the CFS group was not significantly different from the 50% random-response level (*t* (85) = -1.21, *P* = 0.229). The CFS group also showed lower confidence (M = 1.73, SD = 0.65) than the no-CFS group (M = 2.83, SD = 0.08; *t* (95.38) = -15.05, *P* < 0.001).

However, group differences in accuracy and confidence, and even random-level response accuracy, are not sufficient to establish an absence of perceptual awareness in the CFS group. Notably, average confidence of correct responses in this group was low but significantly greater than the minimum value of 1 (*t* (77) = 10.79, *P* < 0.001), suggesting that at least some participants were aware of some of the CSs; learning might thus have arisen from a subset of trials and/or participants where such awareness occurred. To address this, we quantified CS awareness by calculating an awareness index for each participant, ranging in possible values from 0 for no awareness to 1 for full awareness (see Materials and Methods). Although the awareness index of the CFS group (M = 0.28, SD = 0.34) was significantly lower than the no-CFS group’s (M = 0.92, SD = 0.18; *t* (23.93) = -10.19, *P* < 0.001), it was still significantly higher than zero (*t* (85) = 7.59, *P* < 0.001).

Therefore, in order to test our main hypothesis that the reversal of acquired threat responses can be achieved without perceptual awareness, we characterized the quantitative relation between the level of awareness and the magnitude of reversal learning in the CFS group. To control for possible artifacts of regression to the mean (see Supplementary Methods), we followed the recommendation (16) to first examine the correlation between two independent estimates of the awareness index, one calculated from even-numbered trials, the other from odd-numbered trials. Because noise at the measurement level might occasionally yield extreme (i.e., very low or very high) awareness index scores, an association of such randomly-extreme scores with reversal learning (specifically, low awareness with intact learning) could be an artifact. However, it is highly unlikely that across participants, random noise would yield consistent (and similarly extreme) measurements in separate estimates. Due to regression to the mean, if random extreme values occur in one of the two estimates, they are less likely to occur in the other, resulting in a considerable attenuation of any correlation between the two. We found, however, that the two measures were strongly correlated (*r* (84) = 0.96, *P* < 0.001; Figure 3A); participants’ awareness level in one set of trials was overwhelmingly predictive of their awareness in the other set, confirming their reliability as estimates of awareness.

**Figure 3:**
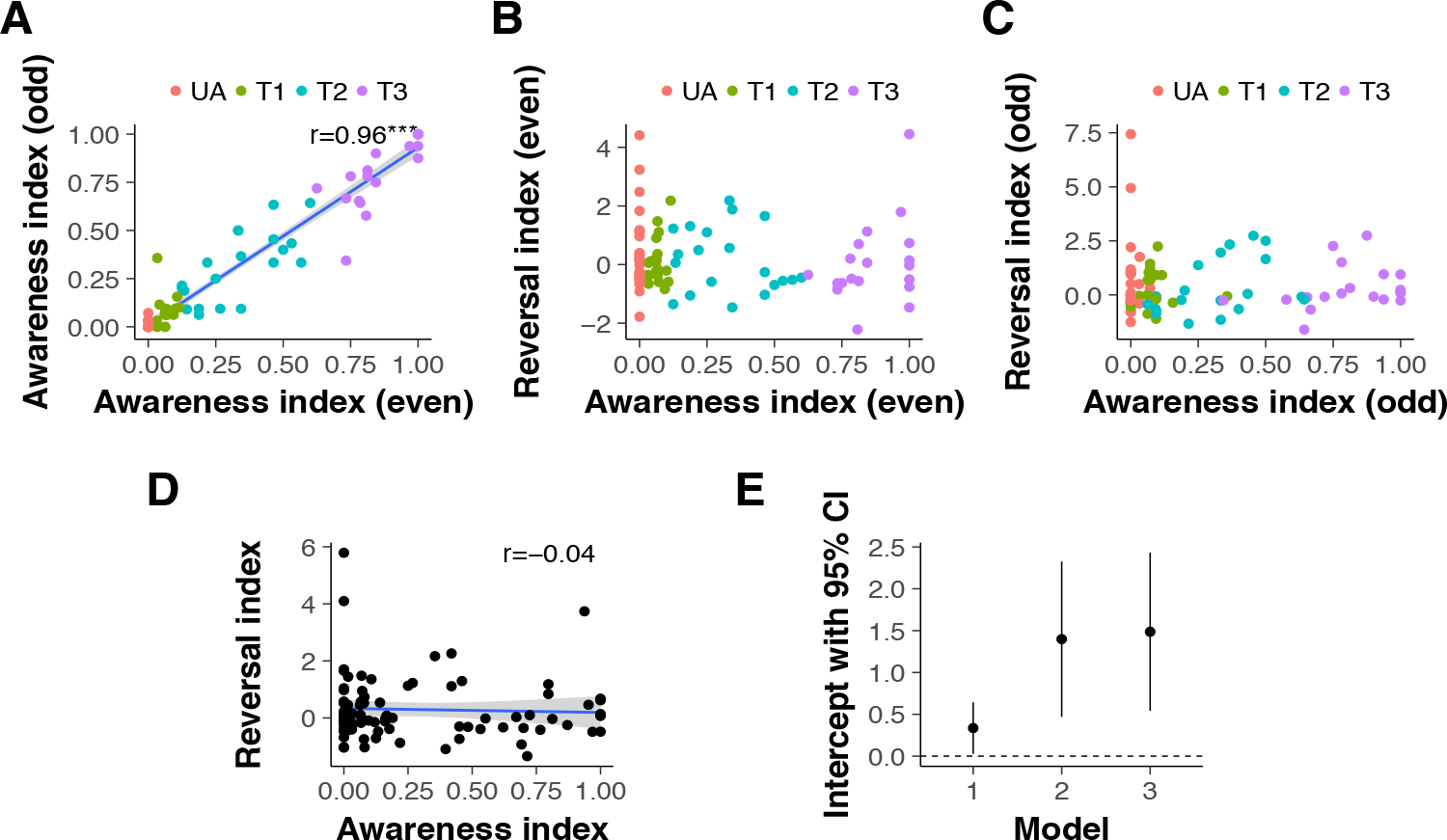
Characterizing the relation between perceptual awareness and reversal learning in the CFS group. **A. Correlation between the awareness index of even and odd-numbered trials**. Each data point represents an individual participant. The strong positive correlation between these independent measures of awareness demonstrates that individual participants’ awareness ratings - even those with extreme values of zero or one - are unlikely to be due to measurement noise. For illustrative purposes, the color scheme marks all participants with an awareness index of 0 in even trials in red (UA, unaware, *N* = 27) and classifies the rest of the CFS group in 3 tertiles (T1-T3). Note that some data points overlap. **B. Reversal learning plotted against perceptual awareness for individual participants, for data obtained from even-numbered trials**. The color scheme is the same as in Panel A. **C. Reversal learning plotted against perceptual awareness for individual participants, for data obtained from odd-numbered trials**. Individual participants are marked with the same color as in the previous panels; the overall distribution of participants is highly similar across panels. **D. Reversal learning as a function of perceptual awareness in the CFS group, using data pooled from all trials**. The intercept, indicating the magnitude of reversal learning in the absence of awareness, is positive and significantly different from zero. **E. Reversal Index intercepts and their 95% confidence intervals in a series of regression models**. Model 1 depicts the intercept (the value of the reversal index when the awareness index equals zero) shown in Panel D. Model 2 shows the intercept when the regression model includes STAIT scores in addition to the perceptual awareness index. Model 3 regresses the reversal index onto the perceptual awareness index, STAIT and tracking scores. (Excluding the potential outlier in the top left corner of panel D weakens significance of the intercept in model 1, *P* = 0.07; the intercepts of model 2 and 3 remain significant after removal of this outlier). Blue lines show linear fits, and ribbons around lines indicate bootstrapped 95% confidence intervals around the estimate.

Next, we examined the association between the awareness index and the reversal learning index, using values of both indices obtained separately from even (Figure 3B) and odd (Figure 3C) trials. As the color-coding of Figure 3 shows, the relation between individual participants’ awareness and their reversal learning was highly consistent across these separate measurements. In light of this, we pooled the data from all trials and regressed the reversal learning index on the perceptual awareness index (Figure 3D). The parameter of interest was the intercept, i.e., the magnitude of reversal learning at zero perceptual awareness. The intercept was positive and significantly different from zero. Furthermore, the awareness index regressor did not contribute significantly to prediction of reversal learning; importantly, this finding was even stronger in models that accounted for STAIT scores and a binary factor indicating whether participants were tracking the stimuli with their responses (see Materials and Methods; Figure 3E and Table 1).

**Table 1:**
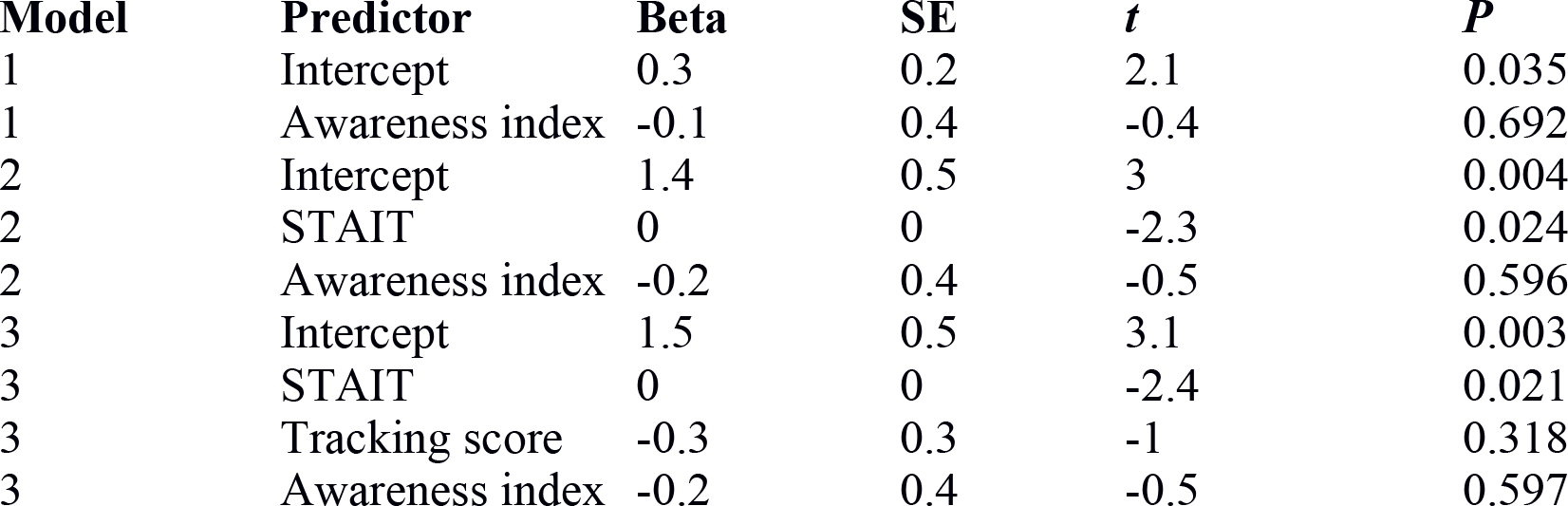
Regression coefficients for all awareness index models. Reversal learning was the dependent variable in all models. Model 1 included an intercept and the perceptual awareness index; model 2 additionally included STAIT scores; model 3 additionally included STAIT and tracking scores.

### Comparing learning and expectation-based accounts

Well-controlled lab-based conditioning procedures require strict constraints that preclude complete randomization of the number and order of different CSs; this comes with a cost: participants are able to develop expectations with above-chance validity, based on the sequence of trials so far, about the likelihood of a shock on any upcoming trial (18). Even without any awareness of the CSs, a participant should have been able to distinguish two types of trials: reinforced (with shock) and non-reinforced (no-shock). In a study with two CSs and a 100% reinforcement rate like ours, such expectations would correspond to an anticipated pattern of alternating trial-types (shock/no-shock or vice versa), with an increase in shock anticipation after every no-shock trial. The question, therefore, was whether the physiological responses we had measured might simply reflect participants’ pattern-based anticipation of shock, rather than learning of the contingencies associated with the CSs.

To answer this question, we used a Bayesian approach to compare the probability of our findings being accounted for by a classic Rescorla-Wagner learning model (22) and a trial-sequence model. We hypothesized that successful threat reversal without perceptual awareness should be better explained by the Rescorla-Wagner learning model, whereas simple pattern-based expectation would be better explained by the trial-sequence learning model. We used maximum likelihood estimation to assess the log likelihood and calculate the Bayesian Information Criterion (BIC) of each model (See Materials and Methods for details of each model and calculation of the BIC). A smaller BIC indicates a better model, and BIC values can thus be compared by calculating the difference between them and interpreting the resulting Δ BIC as providing evidence against the higher BIC. The Rescorla-Wagner model (BIC: 562.1) outperformed the pattern-based expectation model (BIC: 584.9), with the difference (Δ BIC: 22.9) greater than 10, suggesting that the evidence against the trial switch model is very strong (23). Repeating this comparison for just the participants with zero mean awareness confirmed the lower BIC for the Rescorla-Wagner model (BIC: 114.3) compared to the pattern-based expectation model (BIC: 125.7), with the difference again greater than 10 (Δ BIC: 11.3; see also Figure S2). This model comparison provides convincing evidence that a classical Rescorla-Wagner learning model explains our findings better than an alternative expectation-based model.

## Discussion

These results indicate that participants updated their defensive physiological responses independently of their awareness of threat-related cues. Previous studies have shown that new threat associations can be formed without perceptual awareness of the conditioned stimuli (5, 9–10). However, until now it was unknown whether the far more complex process of threat reversal - shifting reactions from a stimulus that no longer predicts danger to one that now does - can be accomplished without awareness. The independence of reversal learning from perceptual awareness, reported here, suggests that separate processes underlie affective flexibility and conscious processing (24). Conversely, the negative correlation between reversal learning and anxiety suggests that the various impairments caused by anxiety are not limited to the systems underlying conscious processes.

Previous studies have pointed out the limitations of using accuracy and confidence measures to assess perceptual awareness, and suggested remedies including the calculation of metacognitive sensitivity measures (25), Bayesian statistics (26), or parametric variation of the experimental manipulation (27). The present study addresses an issue not covered in previous discussions, by showing that a trial-wise analysis may reveal hints for incomplete suppression that analyses relying on average measures might easily miss. Future studies that rely on forced-choice questions for awareness assessment should thus examine response patterns across trials in addition to collecting aggregate measures.

Notably, a previous study (10) that used CFS to investigate acquisition of threat responses without awareness of the stimuli found that such acquisition can occur, but is rapidly forgotten. The present study again showed that such acquisition can occur (and, additionally, be reversed), but did not find the same rapid forgetting. The reasons for this are unclear, but we speculate that the difference may be due to specific aspects of the stimuli, design and procedure: our use of pictures of spiders (rather than the faces used in the previous study) and a 100% (rather than 50%) reinforcement protocol may have altered the temporal characteristics of acquisition. Similarly, the temporal profile of reversal may change if the stimuli and reinforcement regime are different.

The present results add to a growing body of findings distinguishing functions that do and do not require awareness. Such distinctions are important in guiding research into the neural mechanisms of conscious and non-conscious processing. Previous research hints at the mechanism underlying the non-conscious affective flexibility reported here, although it remains to be elucidated: The ability to reverse conditioned responses depends on the integrity of circuitry spanning several neural regions, particularly the ventromedial prefrontal cortex (vmPFC) and its connections with the amygdala (1) where threat associations are formed (28). Consistent with this, it is known that patients with anxiety disorders often show rigid and inflexible threat responses in conjunction with prefrontal cortex dysfunction (29, 30).

Indeed, the real-life settings that people with anxiety disorders find challenging often require the updating and shifting of threat responses. Deficits in affective flexibility may thus explain the threat learning and extinction deficits seen in such disorders (31): Compared to healthy controls, patients are less able to distinguish between safe and unsafe stimuli in threat learning (when it is adaptive to do so), and distinguish between them to a greater extent during extinction (when it is non-adaptive). Threat learning without perceptual awareness is also negatively correlated with baseline state anxiety in healthy participants (10). Our new finding that baseline anxiety is negatively correlated with affective flexibility suggests a potential use for reversal learning as a model paradigm for investigating how anxiety modulates various processes in a variety of disorders, including, for example, posttraumatic stress disorder, in which there is an impairment of threat inhibition (32).

## Methods

### Participants

Ninety-eight healthy participants (mean age = 29.97; range 18-65) were assigned to one of the two groups: reversal learning with CFS (CFS group; *N* = 86, 48 female) or without CFS (no-CFS group; *N* = 12, 5 female). Assignment was random until each group reached a size of 12; subsequent participants were assigned to the CFS group. Measures of trait and state anxiety (Spielberger Trait-State Anxiety Inventory (33); STAIT and STAIS, respectively) and spider phobia (Fear of Spider Questionnaire; FSQ (34)) were taken prior to participation and did not differ between the groups (Table S1). The experiment was approved by the Institutional Review Board of the Icahn School of Medicine at Mount Sinai. All participants provided written informed consent and were financially compensated for their participation.

### Experimental procedure

Participants viewed the stimuli monocularly, through a mirror stereoscope (StereoAids, Australia) placed at a distance of 45 cm from a 17-inch Dell monitor. The CSs (schematic low-contrast images of spiders), presented to the left eye only, were suppressed from awareness in the CFS group: while the left eye saw them, the right eye was presented with “Mondrians” - arrays of high contrast, multi-colored, randomly generated rectangles alternating at 10 Hz. Both the CSs and the CFS masks were flanked by identical textured black and white bars, to facilitate stable ocular vergence. The no-CFS group viewed identical CSs (also presented monocularly), but with no Mondrians presented to the other eye.

The experiment consisted of 16 acquisition trials followed by 16 reversal trials. One of two spider images was presented on each trial. The spider images were schematic and had similar low-level features. During acquisition, spider A always terminated with a shock and spider B never did. Reversal occurred halfway through the experiment: spider B now terminated with a shock and spider A did not. The spider stimuli were presented for 6 s each in pseudorandomized order. One of four possible trial orders was used for each participant. Orders were generated by imposing specific constraints on the trial order, such that the first trial was always reinforced and no more than two of the same trial type ever occurred consecutively.

Trial order and spider identity were counterbalanced across participants. To assess the effectiveness of the awareness manipulation (35), 1 s after the offset of every CS participants were shown the question “Which seen?” (1 = flower, 2 = spider; notably, flowers were never shown, meaning the question addressed detection rather than discrimination as it could be answered correctly even with a brief glimpse). This was followed by the question “How confident?” (1 = guess to 3 = sure; participants were instructed to indicate how confident they were of the flower/spider answer they had just given). Both questions were presented binocularly (1.5 - 2 s each, during which responses had to be given by pressing number keys on a standard keyboard). The second question was followed by an 8 to 10 s inter-trial interval.

### Psychophysiological stimulation and measurement

Mild electric shocks were delivered using a Grass Medical Instruments SD9 stimulator and stimulating bar electrode attached to the participant’s right wrist. Shocks (200 ms; 50 pulse/s) were delivered at a level determined individually by each participant as “uncomfortable but not painful” (maximum of 60 V), during a work-up procedure prior to the experiment.

Skin conductance responses (SCR) were measured with Ag-AgCl electrodes, filled with standard isotonic NaCl electrolyte gel, and attached to the middle phalanges of the second and third fingers of the left hand. SCR signals were sampled continuously at a rate of 200 Hz, amplified and recorded with a MP150 BIOPAC Systems skin conductance module connected to a PC.

### Analysis of physiological responses

#### Model-based analysis

We estimated SNA from SCR data with a model-based variational Bayes approximation (19), inverting a forward model that describes how (hidden) SNA translates into (observable) SCR. A unit increase in SNA corresponds to an increase in SCR of 1 micro Siemens. The model assumes that the observed SCR can be decomposed into different components including anticipation, evocation, and spontaneous fluctuations, each of which are generated by bursts of SNA driven by changes in sympathetic arousal. The generative (forward) model thus describes how sympathetic arousal, the physiological measure that is taken as an index of the psychological process of threat, translates into sudomotor nerve bursts which then generate the observable SCR (19).

Using Bayesian inference, the forward model can then be reversed in order to estimate the most likely underlying SNA given the observed SCR:

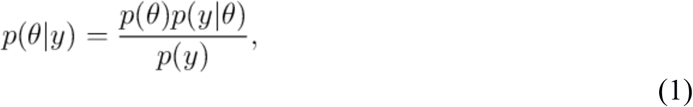

where the most likely parameter vector Θ (corresponding to the SNA) given the observed outcome *y* (corresponding to the SCR) is given by the prior estimate of Θ weighted by the likelihood of *y* given Θ. Solving this equation involves integration over the model evidence p(y) which is analytically hard to compute (and possibly intractable). This can be resolved by replacing this integration problem by an optimization problem, which can be approximated with Variational Bayes procedures (36), where the log of the model evidence can be framed as the sum of the Kullback-Leibler divergence and the Free Energy. By maximizing the Free Energy the Kullback-Leibler divergence is minimized, and a lower bound to the log model evidence can be derived iteratively.

The SNA estimates were computed using previously developed software package PsPM (19) implemented in MATLAB R2016b (The Mathworks Inc, Natick, MA, USA). The statistical analyses were conducted with the R software (37) (R version 3.4.2 (2017-09-28)) and the libraries lme4 (38) and lsmeans (39). Welch’s t-tests were used instead of two sample t-tests when groups had unequal variances.

#### Reversal Learning Index

An estimate of SNA was obtained for each trial. We expected Spider A to evoke greater SNA than Spider B during the acquisition phase, and Spider B to evoke greater SNA than Spider A during the reversal phase. The strength of reversal learning can thus be quantified by calculating, separately for the acquisition and reversal phases, the difference between the average SNA evoked by each spider. To quantify the degree of reversal (which is formally equivalent to the interaction of phase and stimulus), the reversal learning index was calculated by subtracting the difference between mean SNAs evoked by each spider during reversal from the difference during acquisition (the larger the index, the greater the magnitude of reversal learning):

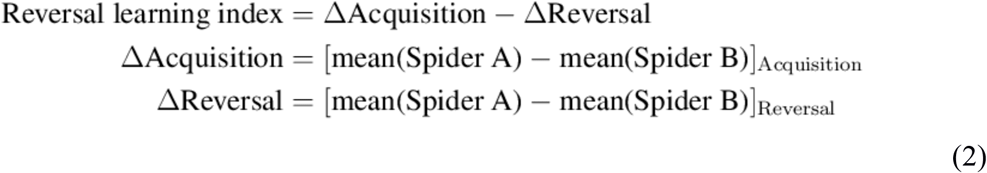

To formally test for group differences in the strength of reversal learning, we computed a linear mixed model using the lme4 library in R. We used the skin conductance response (converted to a model-based measure of sudomotor nerve activity, SNA) as the dependent variable and entered group (CFS, no-CFS), stage (acquisition, reversal), and spider (spider A, spider B) as well as a continuous variable for trial (to account for habituation) as predictors. The random structure of the model included an intercept and slopes for stage and spider.

### Assessments of perceptual awareness

#### Perceptual awareness index

To characterize participants’ reported awareness of CSs, each trial was assigned a perceptual awareness score, defined by a combination of detection and confidence responses: Correct answers with a confidence rating of 1 (guess) and incorrect answers irrespective of confidence were assigned an awareness score of 0; correct answers with a confidence rating of 2 (medium) were assigned a score of 0.5, and correct answers with a confidence rating of 3 (high) were assigned an awareness score of 1. A perceptual awareness index was calculated for each participant by averaging awareness scores across all trials.

#### Stimulus-response association patterns (“tracking”)

We also assessed response patterns across trials, to see whether participants were able to track stimuli with their responses, accurately discriminating the images despite not being able to label them. We plotted individual trial-by-trial responses to the question “Which seen?”, overlaid on the trial-by-trial presentation of spiders (spider A, spider B; Figure S1A). We then calculated the number of consecutive “hits”, defined as the number of consecutive trials where these two time-courses were either identical or consistently in opposition, suggesting that there was a possible association between the stimulus and the response during those trials. The probability of such consecutive hits occurring by chance alone can be derived as follows:

Let *p* = 0.5 be the probability of a hit, *k* the number of consecutive hits, *n* the number of trials left, *i* the number of consecutive hits already observed; the chance of observing *k* consecutive hits for the remaining *n* trials can then be formulated as a recursive problem:

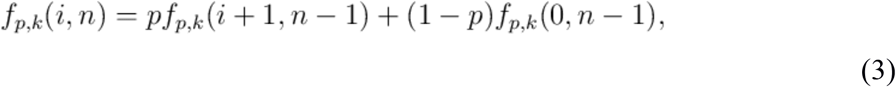

which can be solved analytically with dynamic programming or recursion. Trivially, *f_p_, k*(*k, n*) = 1 for *n* ≥ 0 since *k* consecutive hits have already been observed, and *f_p_, k* (*i, n*) = 0 for *k* – *i* > *n* since there are not enough trials left to observe *k* consecutive hits.

For example, assuming we want to know how likely it is to observe *k* = 8 consecutive hits within *n* = 32 trials given *p* = 0.5, i.e., *f*0.5, 8 (0, 32), we find that this yields a probability of 0.050.

Alternatively, the probability can be derived by simulation for all possible numbers of consecutive hits within 32 trials (i.e., from 1 to 31). For each possible number, we thus also simulated 10^5^ draws of a binomial distribution and calculated the average probability of that number of hits being consecutive. As can be seen in Figure S1B, the result for 8 consecutive hits (0.04991) was very close to the analytical solution. Fifteen participants showed evidence of tracking the spiders or the shocks with their responses (8 or more consecutive hits); notably, 3 of these participants appeared to have a perceptual awareness index of zero. We thus adjusted our subsequent analysis with an additional binary covariate, indicating whether participants did or did not show 8 or more consecutive hits.

### Comparing learning and expectation-based models

The Rescorla-Wagner model (22) describes how the prediction for each trial is updated according to a prediction error and learning rate:

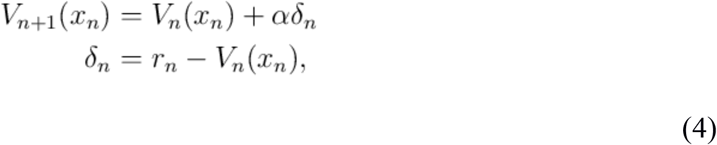

where *x_n_* is the conditioned stimulus on trial *n* (Spider A or Spider B), and δ*_n_* is the punishment prediction error that measures the difference between the expected and the actual shock (*r_n_*) on trial *n*. The learning rate α for the value update is a constant free parameter. The value for the CS not observed on trial *n* remains unchanged. To derive the best fits for the Rescorla-Wagner model, we assumed that *V*_0_ = 0.5, reflecting the assumption that getting a shock or not was equally likely for the first trial.

For the alternative trial-sequence learning model, we assumed that a participant expecting a strict sequence of alternating trial types (shock/no shock or vice versa) would update this expectation according to the actually encountered trial types and a constant learning rate:

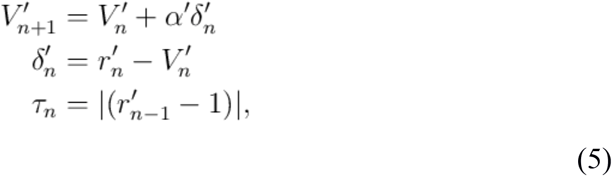

where *V*′_*n*+1_ is the expected trial type switch at trial *n*+1 (if *V*′_*n*+1_ is larger than 0.5, a trial switch is expected), α′ is the learning rate, and δ′_*n*_ is the prediction error. The prediction error corresponds to the difference between the actual trial type switch for trial *n* (*r′_n_*; coded as one for a trial type switch and zero for an equal trial type) and the expectation for trial n. A changing trial type for trial n was tracked by *τ_n_*, which was one if the preceding trial was zero and zero if the preceding trial type was one. To map these expectations onto expected values, we assumed that

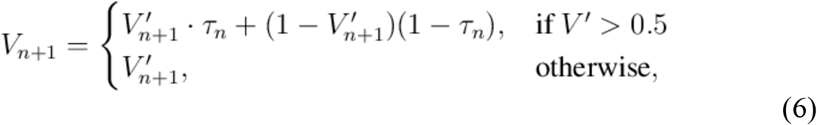

where the expected value for trial *n*+1 was calculated according to whether a trial type switch was expected (*V*′ > 0.5) or not.

We performed a formal model comparison between the conventional Rescorla-Wagner model and the trial switch model for our data set (Figure S2), using maximum likelihood estimation and non-linear optimization (implemented with the fmincon function in MATLAB R2016b (The Mathworks Inc, Natick, MA, USA). Using the log likelihood, we calculated the Bayesian Information Criterion (BIC) to compare the two models as follows:

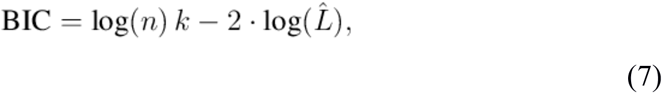

where *n* is the number of data points, *k* is the number of regressors, and 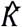 is the maximized value of the likelihood function.

## Acknowledgments

We thank Patrik Vuilleumier who created and shared the spider stimuli. This work was supported in part through the computational resources and staff expertise provided by Scientific Computing at the Icahn School of Medicine at Mount Sinai. Funding was provided by NIMH grant MH105515 and a Klingenstein-Simons Fellowship Award in the Neurosciences to D.S.; ERC Advanced Grant XSPECT-DLV-692739 to D.C. (Co-I); and Swiss National Science Foundation grant SNF 161077 to P.H.

## Author Contributions

PH carried out the computational modeling and statistical analysis, interpreted the results, and drafted the manuscript. HL prepared materials, collected the data and critically revised the manuscript. IL contributed to the conception of the study, the computational modeling, the interpretation of the results, and critically revised the manuscript. CMR contributed to the interpretation of the results and critically revised the manuscript. DRB contributed to the computational modeling and critically revised the manuscript. DS conceived, designed and coordinated the study, contributed to data analysis and interpretation, and critically revised the manuscript. DC contributed to the conception of the study, data analysis and interpretation of results, and drafted the manuscript in its final form. All authors gave final approval for publication.

