## Supplementary Information for "Affective flexibility without perceptual awareness"

### Affective flexibility without perceptual awareness: Supplementary Information

#### Supplementary Methods

Investigating unconscious perception relies on the effectiveness of the technique used to suppress stimuli from awareness. Although CFS is highly effective, it is not foolproof - different observers are not equally susceptible to it, and stimuli often break through the suppression. The standard approach to dealing with breakthrough of the suppressed stimulus is to remove from analysis those trials and individuals in which it occurred. This approach is problematic, though, as it may lead to various artifacts. Below we detail the concerns that are addressed by the analysis approach we adopted in the present study.

##### Regression to the mean

Recent computational work suggests that findings in the trials that remain after removing suppression failures (unless the number of such trials is negligibly small) could be the result of regression to the mean (1): If two noisy measures of the same underlying phenomenon are used (e.g., behavioral and skin conductance responses may both measure conscious processing, each with its own, independent measurement noise), then selecting only the extreme cases of one measure (e.g., only those cases where behavior indicates complete absence of awareness) is unlikely to yield similarly extreme results for the other measure (e.g., it is unlikely that the skin conductance responses will also be close to zero). Thus, a result that looks as if it indicates unconscious processing would in fact be an artifact that is entirely due to regression to the mean.

To address this issue, we refrain from removing trials or participants with breakthrough. Instead, we include all of them in our analyses and assess CS awareness in a continuous manner by assigning each participant an awareness rating; we base this rating on two independent measures of awareness (odd and even-numbered trials), as proposed by Shanks (1) to control for regression to the mean in awareness estimates. We then examine the association between the level of CS awareness and the amount of reversal learning indicated by physiological stimulus-evoked responses (Figure S1).

#### Sensitivity of awareness assessments

The tasks used to assess whether participants were aware of the suppressed stimulus (typically through examining, after each trial, whether they can confidently and accurately report which stimulus was presented) may not be sensitive enough to detect such awareness. This is because breakthrough, leading to at least some awareness, may not always reach a level that allows above-chance performance on the chosen discrimination task (e.g., one might be able to distinguish between two stimuli without being able to label either, a problem that falls within the general concern of "sensitivity dissociation" between direct and indirect measures; (2)).

To address this issue, we analyze individual participants' response patterns across trials, searching for any link between stimuli and behavioral responses - including consistently wrong ones - that may indicate potential tracking of the stimuli, suggesting some perceptual awareness (even in the absence of an explicit ability to identify the images) and thus possibly accounting for physiological findings. We incorporate a binary factor indicating whether or not tracking behavior was observed in each participant into the regression model accounting for physiological responses.

#### Participants' expectations

Laboratory models of conditioning require a pre-defined number of trials for each CS category, to avoid confounding learning with exposure to different stimulus ratios. Furthermore, the order of trials is often not completely random but constrained to allow for roughly similar distributions of CSs across the experiment (3, 4). This means that participants would not be wrong to predict the probability of receiving aversive stimulation on a given trial using a heuristic that takes into account the overall rate of the aversive stimulus and the number of trials since its last occurrence (this is akin to the gambler's fallacy, but in such cases has a higher-than-chance success rate). The likelihood of participants basing their anticipation of a shock - and of the physiological arousal that comes with such anticipation - on such probabilistic estimates may be even higher when CSs are suppressed from awareness, leaving the overall rate and distribution of shocks as the only bases for such predictions. Indeed, such expectations may account, at least in part, for some of the findings of previous non-conscious learning studies (5). We address this issue by using Bayesian model comparison, demonstrating that a model featuring updating of learning provides a better account for the findings than a shock-expectation model based on the distribution of shocks across the experiment.

### Supplementary Figures

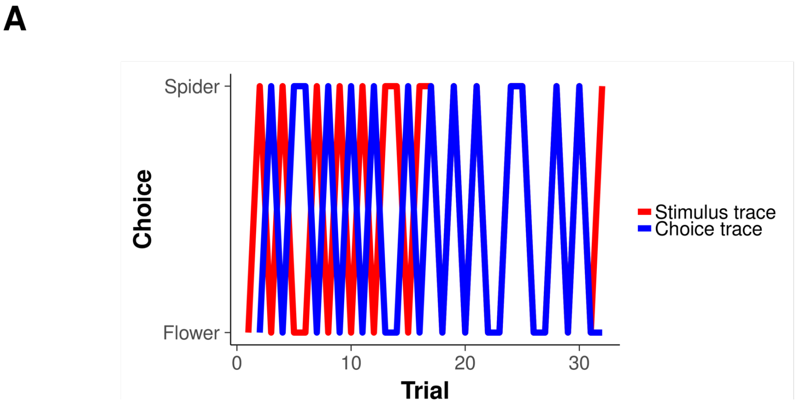


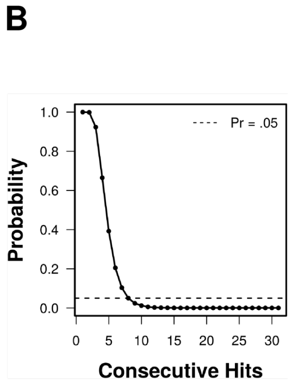


**Figure S1: Examination of tracking behavior in the correspondence between time courses of forced-choice responses and stimulus presentation in the CFS group. A. Forced choice responses (blue) versus stimulus presentation (red) in an illustrative participant.** This participant shows periods in which forced-choice responses track or are in direct opposition to both the presented stimuli and shock application, suggesting that this participant tracked the spider and/or the shock with his responses. **B. Probability of consecutive hits ($10^5$ simulations).** Hits were defined as the number of consecutive trials where the time courses of forced-choice responses and stimulus presentation were either identical or directly opposed to each other. To quantify the probability of consecutive hits, we simulated the probabilities for all possible levels of consecutive hits (e.g., from 1 to 31 for 32 trials). We simulated $10^5$ draws of a binomial distribution, and calculated the average probability to see (at least) each possible number of consecutive hits. The probability exponentially declines as the number of consecutive hits increases, reaching a probability of about 5% at 8 consecutive hits. Thus, observing 8 or more consecutive hits in 32 trials in a perceptually unaware participant who is truly guessing is unlikely, which is why we considered this an indication for stimulus tracking behavior.

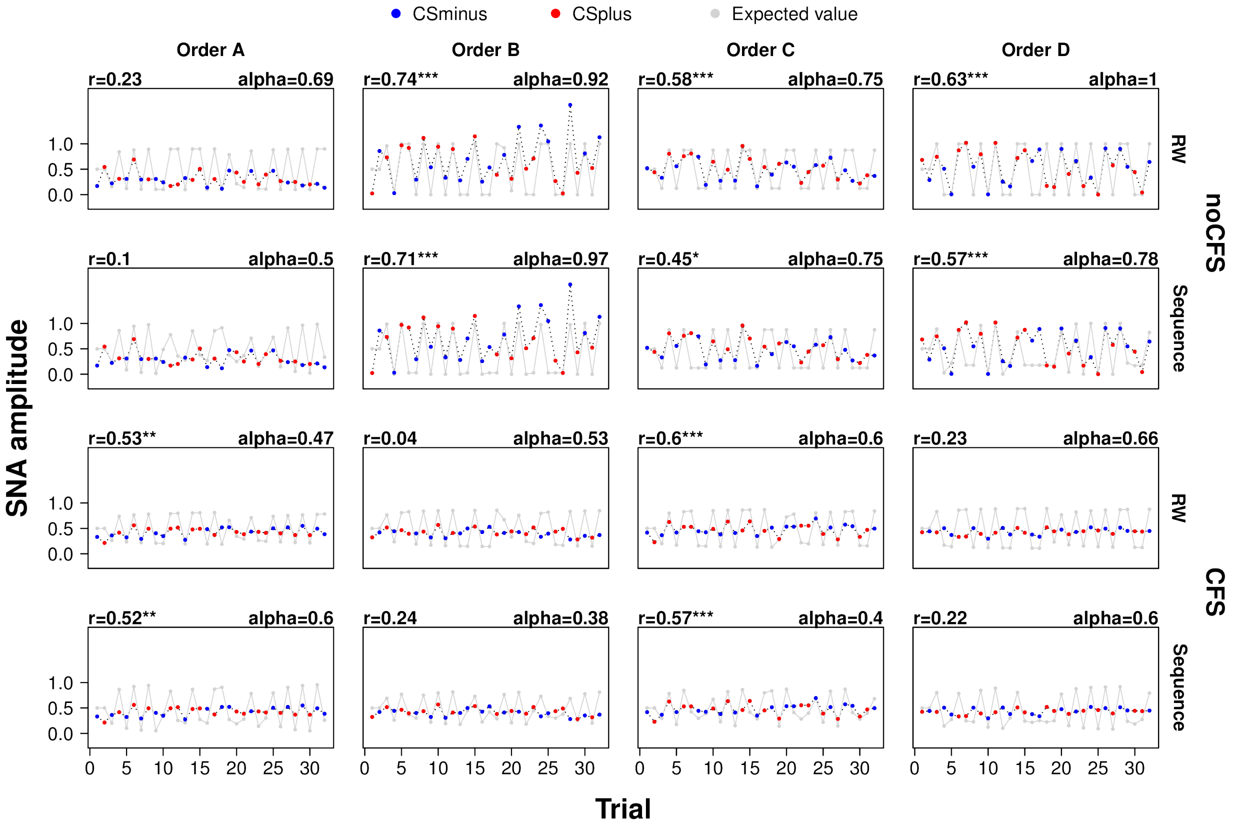


**Figure S2: Time series of measured and predicted threat response indicate that the Rescorla-Wagner learning model explains the data better than an alternative trial-sequence learning model that simply assumed a sequence of alternating trial types (no-shock/shock or vice versa).** Trial-wise mean responses to CS+ and CS- are shown by group, learning model and experimental order. Each panel also shows the Pearson correlation between predicted and measured time series and the learning rate (parameter alpha in each of the learning models). The generally higher Pearson correlation coefficients are consistent with the formal model comparison between the Rescorla-Wagner model and the trial-sequence model, which confirmed that the Rescorla-Wagner model provided stronger evidence than the alternative learning model. r, Pearson correlation coefficient; alpha, learning rate; RW, Rescorla-Wagner learning model; Sequence, trial-sequence learning model; SNA, sudomotor nerve activity; ***, $P < 0.001$; **, $P < 0.01$; *, $P < 0.05$.

#### Supplementary Table

| Characteristic | CFS N | Mean | SD | no-CFS N | Mean | SD |
|---|---|---|---|---|---|---|
| Male | 38 | | | 7 | | |
| Female | 48 | | | 5 | | |
| Age, y | 86 | 29.7 | 8.3 | 12 | 31.7 | 10.6 |
| STAIT | 80 | 34.2 | 9.6 | 11 | 33.5 | 12.3 |
| STAIS | 80 | 29.4 | 8.6 | 11 | 28.4 | 9.2 |
| FSQ | 79 | 43.5 | 21.5 | 12 | 36.8 | 12.1 |
| UR, SNA units | 86 | 1.8 | 0.5 | 12 | 1.8 | 0.7 |

**Table S1: Sample characteristics.** Note: STAI data for 7 participants (6 of them in the CFS condition) and FSQ data for 7 participants (all of them in the CFS condition) were lost due to an archiving error. Exclusion of these participants from subsequent analyses does not alter the overall pattern of results. *Abbreviations*: CFS, Continuous Flash Suppression; SD, Standard deviation; STAIS/STAIT, state/trait anxiety subscale of the Spielberger State-Trait Anxiety Inventory; FSQ, Fear of Spider Questionnaire; UR, unconditioned response (shock response); SNA, sudomotor nerve activity.
